# Biologically-constrained insect models enable realistic simulations and improve biomechanical predictions of locomotion

**DOI:** 10.1101/2025.11.13.687989

**Authors:** Omer Yuval, Elad Ozeri, Avi Amir, Amir Ayali

## Abstract

Animal locomotion emerges from the complex interplay of morphology, neural control, and biomechanics, involving both active and passive elements. Insect have attracted much interest, and a rich ecosystem of computational models exists for insect locomotion. However, while active control mechanisms have been extensively studied, obtaining accurate and useful models remains challenging due to limited understanding of how detailed anatomical constraints and passive biomechanical properties shape movement. To address this, we developed anatomically accurate 3D models of the adult desert locust and mole cricket, two orthopteran insects that share a basic body plan but exhibit strikingly different morphologies linked to their habitats and specialized locomotor behaviors — jumping in locusts versus subterranean digging in mole crickets. We fine-tuned these models using precise morphometric measurements for each major body segment. Furthermore, we quantified passive joint dynamics through high-speed videography of anesthetized locust, revealing a two-phase return motion and history-dependent resting angles. By integrating these biologically grounded constraints into physical simulations, we significantly narrowed the parameter space, resulting in more realistic simulations. Our approach provides new tools for predicting biologically relevant, yet experimentally challenging variables, such as joint torques and contact forces, which contribute directly to the understanding of the underlying biological phenomena. Moreover, these improved simulations offer valuable insight for designing energy-efficient soft robotic systems.

## 1 Introduction

Animal locomotion emerges from the integrated interplay of morphology, neural control, and biomechanics [1]. Making up approximately 75% of all known animal species, insects are the most diverse animal group, employing an extremely wide range of locomotor strategies to walk, jump, climb, dig, and fly. Insect locomotor performance is tightly linked to morphological traits such as segment dimensions, mass distribution, and joint properties, which together determine mechanical leverage, energy storage capacity, and gait stability. Their rigid exoskeleton, which is jointed and multi-segmented, imposes significant mechanical constraints on movement. Passive biomechanical elements — structures and materials that generate forces without active muscle contraction — are thus particularly important in insect locomotion [2–6].

Such passive elements include elastic cuticle structures, resilin pads, compliant joints, and passive muscle properties. Operating within the geometric limits set by limb and body morphology, passive biomechanical properties contribute significantly to energy storage and release during rapid movements, facilitate mechanical stability, and provide swift responses without requiring neural input [7]. For example, elastic cuticular components in locust legs efficiently store and release energy during jumps [3]. Similarly, passive joint torques in stick insects can rival active muscle forces, enabling rapid reflexive responses independent of neural activity [8]. At the same time, whole-body and limb morphology — segment dimensions, mass distribution, and joint geometry — sets the mechanical leverage and kinematic workspace within which these passive elements act, thereby shaping attainable gait timing, stability, and energetic costs. While active control mechanisms have been extensively studied, the role of body morphology alongside passive biomechanical properties in shaping insect locomotion remains much less explored, leaving important knowledge gaps.

A diverse ecosystem of computational models already supports insect locomotion research, spanning reduced-order template dynamics and spring-mass models (e.g., SLIP) to full-body neuromechanical simulations [1, 9–14] and physics-engine implementations [12, 13]. While these models show that core gait phenomena can be reproduced in silico and provide a substrate for hypothesis testing, their accuracy and usefulness hinges on how closely morphology and passive mechanics are constrained by real-world experimental measurements. Oversimplification of such properties risks introducing inaccuracies and potentially misleading results.

In order to demonstrate the advantages of biologically-constrained models in improving biomechanical predictions of locomotion, we developed anatomically-grounded models for two orthopteran insects that represent markedly different ecological niches and locomotion strategies: the desert locust (*Schistocerca gregaria*) and the cricket (*Gryllotalpa spp*.). The former adapted for powerful jumping and long-range walking as part of swarms, while the latter specializes in tunnel navigation and subterranean digging, using robust forelegs. We constructed anatomically accurate 3D models, representing each body part as a distinct segmental mesh using Blender [15]. These models were fine-tuned with empirical morphometric measurements collected from adult individuals of both species. The meshes were then exported into the MuJoCo physical simulation environment [12], where mass was assigned based on biological measurements, and degrees of freedom were added between the different body parts in the form of 1D hinge joints.

To specifically quantify passive joint properties, we experimentally measured the dynamics of the hind femur-tibia joints in anesthetized locusts via high-speed videography, observing a biphasic return motion — a rapid initial phase followed by slower settling — and a history-dependent resting angle. This experiment was then replicated in silico in MuJoCo, where joint parameters were manually adjusted to obtain dynamics similar to the dynamics observed in-vivo.

Finally, to test the behavioral effect of the observed passive joint dynamics, we simulated a walking locust and compared body speed and the energetic cost between the adjusted hind femur-tibia joint model and a less biologically-informed, baseline model. Overall, this approach enables us to systematically evaluate how morphology and passive biomechanical constraints jointly shape insect locomotor dynamics.

Accordingly, the study proceeds in three parts: (i) morphometric measurements and construction anatomically grounded models, (ii) comparative analysis of learned double-tripod coordination across insect species, and (iii) experimental-computational mapping of passive hind FT mechanics and their dynamical consequences for simulated locomotion.

## 2 Results

### Functional leg adaptations are manifested in both shape and mass

We constructed segmented 3D models of the desert locust (Fig. 1A) and mole cricket (Fig. 1B) in Blender, guided by 2D projections of each insect’s morphology (Fig. 1C(i-ii)-D(i-ii)).

**Figure 1.**
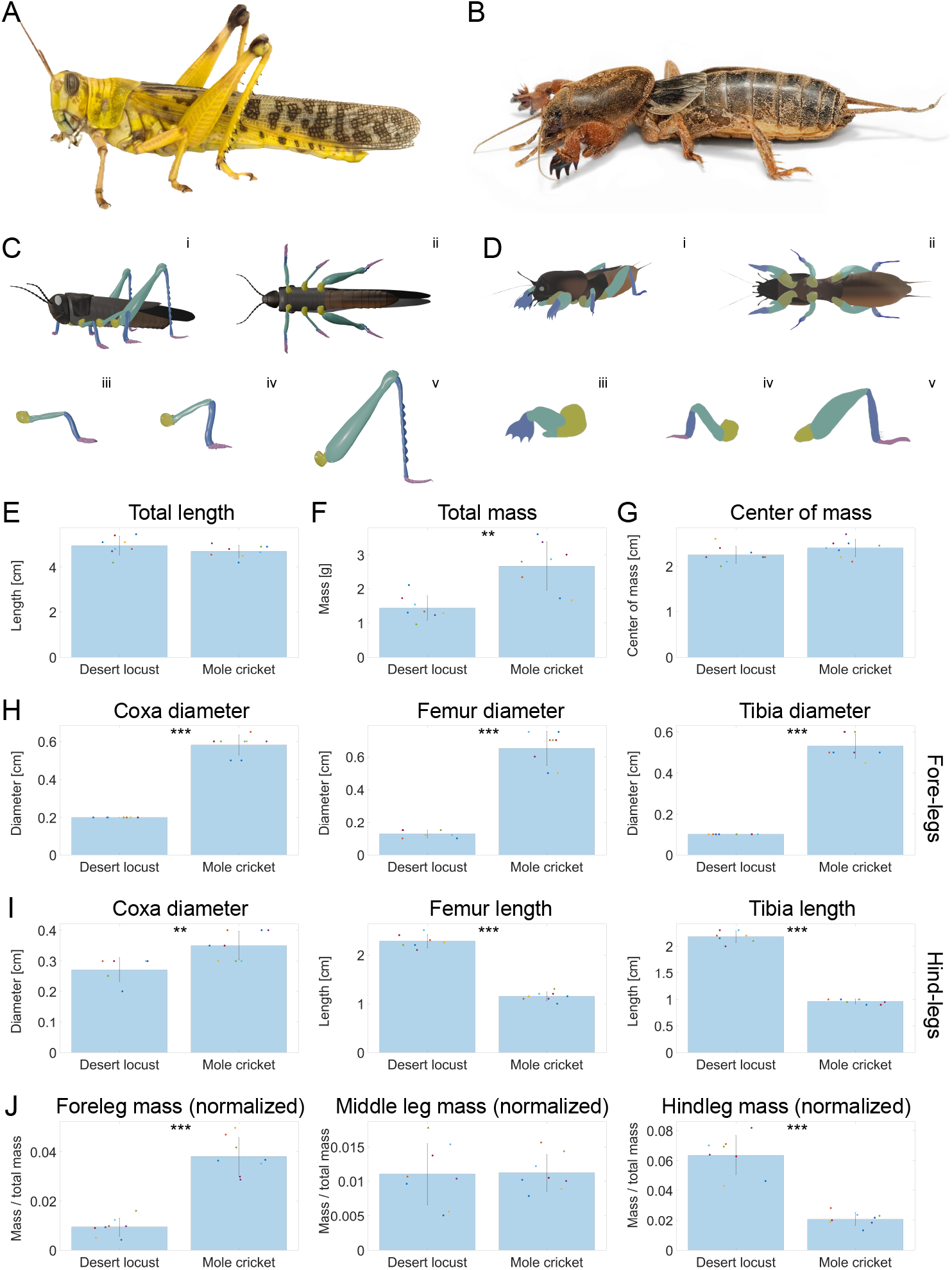
A. Constructing anatomically-grounded insect models. A-B. The desert locust (A) and mole cricket (B). C-D. Models of the desert locust (C) and mole cricket (D). Panels iii-v show the foreleg, middle leg and hindleg, respectively. The models are bilaterally symmetrical. E-G. Total length, measured from the head tip to the abdomen tip (E), total mass (F) and center of mass, measured as longitudinal distance from the tip of the head (G). H. Diameter of the foreleg coxa, femur and tibia. I. Diameter of the hind coxa and length of the hind femur and tibia. J. Total leg mass of the fore, middle and hind legs divided by total body mass.

In order to capture insect-specific anatomy, we measured their dimensions, mass, and center of mass across eight male and female adults per species. Each model was segmented into the head, thorax, abdomen, and leg segments — coxa, femur, tibia, tarsus (Fig. 1C(iii-v)-D(iii-v)). The empirical data guided iterative refinement of the segmented meshes in Blender to ensure faithful qualitative and quantitative representation of shape and proportions.

To assess how overall body proportions and limb specializations reflect each species’ locomotor niche, we compared several key morphological properties between desert locusts and mole crickets. First, we compared total body length (Fig. 1E) to test whether the two species maintain conserved longitudinal dimensions despite divergent lifestyles. Contrary to our expectation, both species exhibited nearly identical mean body lengths. The mole crickets are significantly heavier, consistent with their heavily sclerotized, digging-adapted exoskeleton (Fig. 1F). The longitudinal center of mass, measured as longitudinal distance from the tip of the head, was found to fall at a similar position in both species (Fig. 1G).

As hypothesized, mole crickets showed significantly greater diameters in all foreleg segments — reflecting their role in substrate displacement (Fig. 1H) — while locusts possessed hind-leg femora and tibiae approximately twice as long (Fig. 1I), supporting elastic energy storage and high-power jumping. Conversely, the diameters of hind-leg segments were larger in mole crickets, most pronounced in the tibia (~ 0.1 cm in desert locusts vs. ~ 0.2 cm in mole crickets).

By normalizing each leg’s combined mass (coxa + femur + tibia) to the total body mass (Fig. 1J), we confirmed that the forelegs account for a larger proportion of body mass in mole crickets, whereas locusts allocate more mass to their hind legs, underscoring the functional trade-offs in limb investment between digging and jumping adaptations.

In summary, although mole crickets exhibit significantly greater total mass (Fig. 1F) and pronounced foreleg hypertrophy (Fig. 1H), and locusts allocate more mass to their hind legs (Fig. 1I-J), the longitudinal center of mass remains nearly identical in both species (Fig. 1G). This invariance indicates that the localized mass increases in specific limbs are offset by corresponding decreases or redistribution of mass in other segments, thereby preserving overall balance. Such counterbalancing of segmental masses allows each insect to specialize — jumping in locusts and digging in mole crickets — without compromising postural stability.

### Convergent and divergent gait properties in anatomically distinct insects

Given the interspecific differences in body morphology and mass distribution (Fig. 1), we asked which aspects of tripod walking converge across species and which diverge due to anatomical constraints.

To investigate the mechanical and dynamic consequences of distinct morphologies and passive mechanisms, we exported the refined meshes to MuJoCo, a high-fidelity rigid-body physics engine [12], where the segments were assigned their measured mass and degrees of freedom were added between segments (see Methods).

To this end, we trained a reinforcement learning agent using the desert locust and mole cricket models to perform forward double-tripod walking using the same objective, control architecture, environment, and hyperparameters, and then compared gait kinematics and dynamics (Fig. 2A-B and Videos S2 and S3). The objective encouraged forward velocity and anti-phase tripod contact with the ground (see Methods). Apart from body morphology, the models differed only in the empirically measured mass of their body parts, and in the angular range of their joints, based on previous empirically measured values during double-tripod walking in these insects [14, 16]. Active control was scaled in proportion to each insect’s total body mass by scaling the gear ratio of the motor associated with each joint; all other parameters were identical.

**Figure 2.**
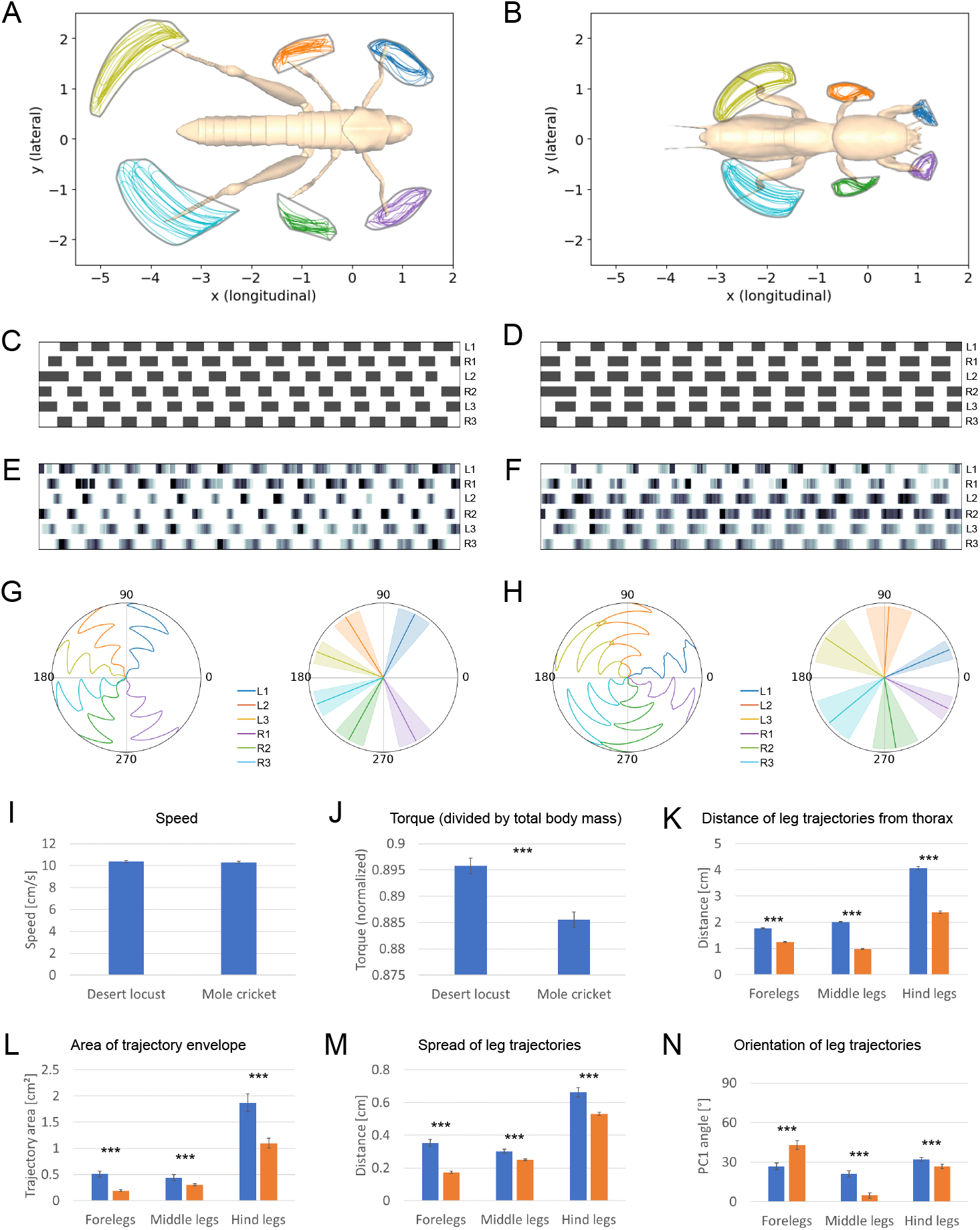
Comparison of simulated gait parameters between the desert locust and mole cricket. A-B. Leg tip trajectories during double-tripod forward locomotion of the desert locust (A) and mole cricket (B) from a top view (left) and side view (right). C-D. Swing-stance diagrams based on leg tip position relative to the body position and orientation (longitudinal body axis). Dark rectangles indicate stance. E-F. Swingstance diagrams based on contact with the ground. Darker color indicates stronger contact force. G-H. Leg angles over time (G; time is shown in the radial direction) and mean*±*std of leg angles statistics (H; see Methods). n=20 simulations. I-J. mean body speed (I) and mean joint torque divided by total body mass (J). In J, the y-axis is truncated above zero; values should be interpreted accordingly. K. The distance of each leg trajectory’s mean coordinate from the thorax coordinate (see Methods). L. Area of the envelopes (gray contours in A-B) of leg trajectories (see Methods). M. Spread of leg trajectories around their mean coordinate (see Methods). N. Orientation of leg trajectories (see Methods). In K-N, values of left and right leg pairs were averaged. n=20 simulations. \*\*\**p <* 0.001.

Swing-stance rasters derived from leg-tip kinematics (Fig. 2C-D) and from ground-contact forces (Fig. 2E-F) both reveal robust double-tripod coordination, as expected given the reward component that minimizes the contact time within each tripod, and enforces an inter-tripod delay of half the target period (see Methods). However, subtle differences are visible here too — in the mole cricket, legs within each tripod are in-phase, while in the desert locust there is a small shift (e.g., between R2 and L3). Moreover, ground-contact forces show that the middle legs in the desert locust exert less ground force and less uniformly, while in the mole cricket the same is true for the forelegs (Fig. 2E-F).

Limb orientation differed systematically between species (Fig. 2G-H). Although both models have sagittal-plane joint limits exceeding 100^*◦*^, the middle legs in each species operated within markedly smaller and non-overlapping angle ranges, indicating convergence to species-specific operating windows. This reduction in used range, despite generous anatomical limits, suggests that the learned controllers exploit morphology — and the soft nature of the objective — to prioritize forward progress and stability rather than using the full permissible kinematic space.

To assess gait efficiency, we compared whole body speed and joint torques (normalized to total body mass). This revealed an unexpected result: both models achieved the same speed (Fig. 2I), but the mole cricket did so more efficiently (Fig. 2J).

A detailed analysis of leg trajectories revealed systematic differences, manifested in their distribution and orientation, despite similar overall path shapes. First, in the mole cricket leg trajectories were significantly closer to the body (Fig. 2K). Second, all leg trajectories were distributed over a larger area in the desert locust, indicated both by the distance from the mean trajectory coordinate and by the area enclosed within its bounding envelope (Fig. 2L-M). Finally, the trajectories of the mole cricket’s middle legs were closer to 90^*◦*^ relative to the main body axis, while the opposite was true in the forelegs. The orientation of hind leg trajectories was similar in both species, but in the mole cricket they were slightly more aligned with the main body axis (Fig. 2N). Expectedly, the orientation of the trajectories of the forelegs and middle legs are in line with leg angle statistics (Fig. 2G-H), as both capture their range of motion.

Overall, both models converged on stable double-tripod coordination, yet they diverged markedly in segment kinematics and propulsive efficiency at comparable forward speeds. By anchoring control to measured body parameters, we link morphometric differences to the observed kinematic patterns and open a path to mechanistic insight into how body form shapes gait and cost of transport.

### Passive joint dynamics shows a biphasic trajectory and history-dependent resting states

Passive joint mechanics play a critical role in insect locomotion, contributing to limb stabilization, energy storage, and rapid recovery from perturbations [3, 7, 8, 17, 18]. To establish a baseline understanding of these passive properties and to further exemplify the great importance of real-world constraints on the performance of computer models, we chose to focus on a specific example — the passive joint characteristics of the femur-tibia joint of the locust hind legs.

We asked how quickly the passive FT joint returns to equilibrium after perturbation, and whether return rate scales with displacement (Fig. 3A-C). Across a range of initial flexion and extension angles, the tibia exhibited a clear biphasic return—an initial rapid movement followed by slower settling (Fig. 3D). The peak initial angular speed (Fig. 3E) increased approximately linearly with the magnitude of the perturbation in both directions (Fig. 3F; extension fit *R*^2^ = 0.81, flexion *R*^2^ = 0.34), with the weaker flexion fit likely reflecting its smaller sampled range. These results suggest that the rate of the fast-phase scales linearly with the magnitude of the displacement in both flexion and extension (see Discussion).

**Figure 3.**
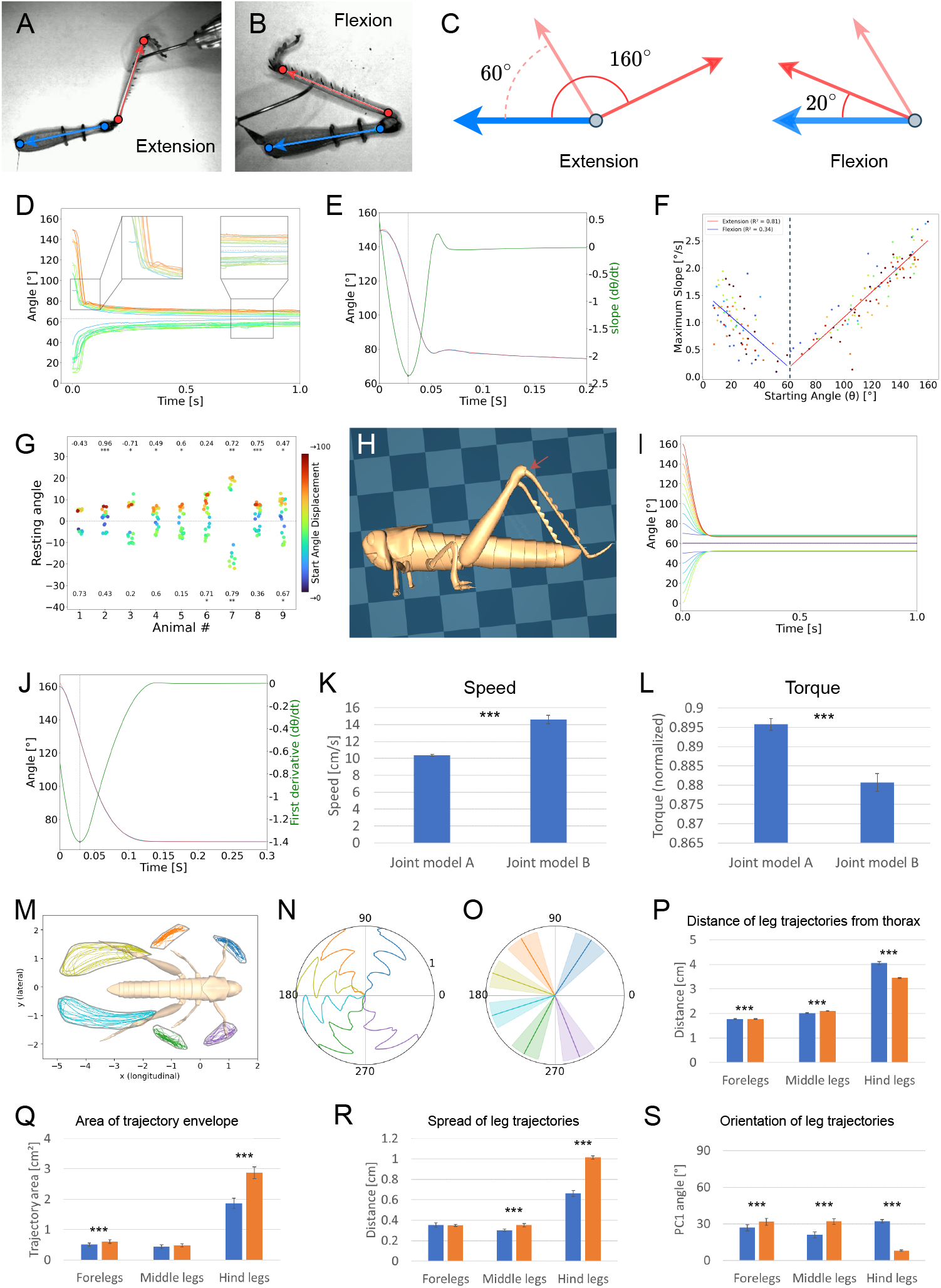
Passive dynamics of the hind femur-tibia (FT) joint in the desert locust. A-B. Extension (A) and flexion (B) poses of the hind FT joint. The angle of the FT joint is measured by tracking two points on the femur (blue) and two points on the tibia (red). C. A schematics of the most extreme extension and flexion displacements (~ 160^*◦*^ and ~ 20^*◦*^, respectively). The semi-transparent arrow shows the resting state (typically ~ 60^*◦*^). D. Angle over time following displacement and release of the tibia. The color corresponds to the magnitude of displacement angle. The left inset shows a rebound reaction, most pronounced following higher magnitude displacements. The right inset shows the history-dependent resting states. E. Time derivative (green, right y-axis) of the most extreme extension shown in D (purple, left y-axis). The local minimum point captures the highest absolute rate of the fast phase, while the local maximum point captures the rebound. F. Maximum slope as a function of initial angle for all animals and initial angles, with linear fits for flexions (blue) and extensions (red). G. Resting angles of each animal. Each dot represents one trial, and its color corresponds to the magnitude of the initial angle. The numbers and stars at the top and bottom show the tau and p-value of the Kendall’s Tau rank correlation coefficient test, indicating how much resting angles are ordered according to their corresponding initial angles. H. A model of the locust’s hind leg. The parameters of the femur-tibia joint were manually adjusted to obtain dynamics similar to the ones observed in-vivo. I. Simulated traces of angle over time (cf. D). J. Time derivative of the most extreme extension in I (cf. E). K-L. Mean joint torque (K) and whole body speed (L) extracted from walking simulations of the locust trained with either the new (model A) or baseline (model B) joint model. M-O. Leg tip trajectories (M) and leg angles (N-O) using the biologically-grounded hind femur-tibia model (cf. Fig. 2A,G). P. The distance of each leg trajectory’s mean coordinate from the thorax coordinate. Q. Area of the envelopes (gray contours in M) of leg trajectories. R. Spread of leg trajectories around their mean coordinate. S. Orientation of leg trajectories. In P-S, values of left and right leg pairs were averaged. \*\*\**p <* 0.001. n=20 simulations. Error bars show standard deviation. In panel L, the y-axis is truncated above zero; values should be interpreted accordingly.

We next tested whether the passive equilibrium is history-dependent. Resting angle (1 s post-release) varied widely across animals and depended on both the magnitude and the direction of the initial displacement (Fig. 3G). In several individuals, larger absolute perturbations yielded larger absolute shifts of the resting angle (strong positive monotonic association; Kendall’s *τ, p <* 0.05), whereas others showed weak or nonsignificant associations; one animal showed a significant negative association (animal #3; Fig. 3G). Positive associations were more common after extensions, plausibly reflecting their larger sampled range. Together, these data demonstrate a history-dependent equilibrium with substantial inter-individual variability.

We also observed a transient “rebound”—a brief overshoot between the fast and slow phases—that increased with perturbation magnitude (Fig. 3D-E), consistent with elastic recoil.

To capture the observed dynamics in silico and test their effects on whole-body locomotion, we replicated the joint experiment in silico. We tuned only the FT passive parameters to match the in-vivo return trajectories. The simulated angle-time traces reproduce the biphasic return and history-dependent equilibria (Fig. 3I) but not the transient rebound (no peak in the angle-derivative for the largest displacement; Fig. 3J).

We then asked whether incorporating the measured FT passive dynamics alters whole-body locomotion. We modified only the hind FT joints: passive parameters tuned to reproduce the in-vivo dynamics, and actuators were scaled so that the stiffer joints remained drivable. Under otherwise identical training and model settings, the biologically constrained model achieved higher forward speed with lower joint torque than the baseline (Fig. 3K-L). This combination of greater speed at reduced effort highlights a direct contribution of passive joint mechanics to both performance and efficiency.

To relate the efficiency gain to whole-body kinematics, we compared the constrained and baseline locust models across leg kinematic descriptors (Fig. 3M-O and Video S3). Both retained double-tripod coordination, but the constrained model reorganized limb use: hind leg tips operated closer to the thorax (Fig. 3P) while sweeping larger envelopes (Fig. 3Q-R), and their paths became more aligned with the body axis; by contrast, fore- and mid-leg trajectories shifted toward ~ 90^*◦*^ (more lateral) orientations (Fig. 3S). Taken together, these changes indicate a redistribution of workspace across the limbs induced by tuning hind FT passive dynamics—the more axially aligned, larger-sweep strokes direct a greater share of stance force into forward motion, providing a parsimonious explanation for the observed increase in speed at reduced joint torque.

## 3 Materials and methods

### Animal maintenance

Desert locusts (*Schistocerca gregaria*) were reared for many consecutive generations in 60 L aluminum cages and under a controlled temperature of 30^*◦*^C, 35-60% humidity, and a 12:12 h light:dark cycle. The locusts were fed daily with wheat seedlings and dry oats.

Mole crickets (*Gryllotalpa sp*.) were collected at different life stages from the field in the countryside around Tel Aviv, Israel. They were maintained at room temperature (20–25^*◦*^C), each in a separate 750 ml glass jar filled with autoclaved plant soil, and fed on flour beetle larvae, grass roots, and slices of carrot. The soil in each jar was moisturized by sprinkling 100 ml of water every three days.

### Measurement of shape, mass and center of mass

To parameterize our 3D models, we quantified segment geometry, mass, and center of mass for head, thorax, abdomen, and leg segments (coxa, femur, tibia; tarsi were too small to measure reliably) in each of eight adults per species. Lengths were measured on a millimeter grid under a stereomicroscope using digital calipers (resolution 0.01 mm, accuracy *±*0.05mm), applying only enough pressure to stabilize the segment without deformation. Masses were recorded on an analytical balance (accuracy 0.0001 g).

Whole-body center of mass (COM) was determined by balancing the intact insect on a vertical bolt fixed to a leveled platform; the distance from the head tip to the balance point was recorded. Total body length (head-tip to abdomen-tip) was measured on the grid. Bodies were then dissected into head, thorax, abdomen, and legs. For head and thorax, we measured mass and three orthogonal dimensions (length, width, height). Abdomen mass and length were measured, and circumferences at the maximum and second-narrowest segments were obtained by fitting a fine metal wire, cutting it, and measuring its length on the grid.

From each individual we dissected one set of fore, middle, and hind legs. Each leg was separated into coxa, femur, tibia, and tarsus. For coxae (assumed roughly spherical) we measured mass and diameter. For femora (front and middle legs) and tibiae we measured mass, segment length on the grid, and diameter with calipers. The hind-leg femur, being markedly non-cylindrical, was further characterized by measuring its maximum width and height (elliptical cross-section assumption).

### Measurement of passive joint dynamics

To characterize passive FT (femur-tibia) joint dynamics, we anesthetized nine adult locusts using CO_2_ and mounted a single (intact) hind leg in a custom holder: the femur was clamped horizontally, and the tibia was free to pivot. Using a fine needle, we manually displaced the tibia to ~ 16 target angles (20^*◦*^-160^*◦*^) and released it, recording the ensuing return motion at 400 fps (Mikrotron MotionBlitz Cube 4; 320*×*504 pixels). Four anatomical landmarks (two on the femur, two on the tibia) were tracked with DeepLabCut [19]. This was used to define their vectors (Fig. 3A-C) and extract joint angle time series (Fig. 3D).

### Mechanical modeling of insects

We built anatomically accurate 3D meshes in Blender 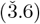 by importing standard 2D anatomical projections and segmenting body parts into separate meshes for the head, thorax, abdomen, and leg segments — coxa, femur, tibia, tarsus (Fig. 1C(iii-v)-D(iii-v)). Quantitative shape adjustments were made by approximating body parts and leg segments with simple 3D primitives (e.g., cuboid, ellipsoid, and cylinder with a variable radius) and fine-tuning the meshes to match measured segment dimensions. The meshes were exported as STL files and imported to MuJoCo (v3.3) [12]. In MuJoCo XML, each segment was assigned its measured mass. Hinge joints were used to add degrees of freedom and were defined as follows: two orthogonal hinges at each coxa-femur junction, one hinge at the femur-tibia junction, and one at the tibia-tarsus junction. Passive joint dynamics was modeled using torsional spring-damper elements, and torque actuators were associated with each hinge for active control.

### Reinforcement learning control and training

We trained locomotion controllers with model-free reinforcement learning (identical settings for both species). The policy output controls joint torques; each action *a*_*t*_ ∈ [−1, 1]^*m*^ (where *m* is the number of actuated DOFs) is linearly scaled by the joint’s gear to produce target torques *τ*_*t*_. Motor gear ratios were scaled in proportion to each model’s total mass. Observations included joint angles/velocities for all actuated DOFs, body orientation, vertical position *z* and linear/angular velocity (horizontal position *x, y* was excluded for translational invariance), and the time since last ground contact for each foot.

Reward function. The total reward was a weighted sum of forward progress, contact-based tripod coordination, orientation and left-right antisymmetry terms, and a constant survival reward. Early episode termination was triggered upon violation of the z-position and orientation thresholds.

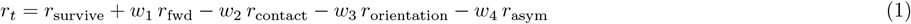

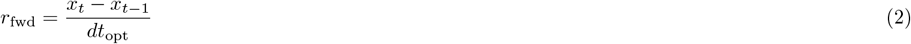

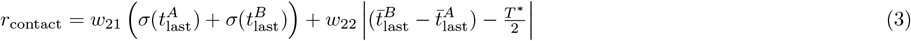

Here *r*_survive_ is a constant; *x* is forward body position; *dt*_opt_ is the optimization time step; for tripod *i* ∈{*A, B*}, 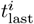 denotes the set of “time-since-last-contact” values for the three legs in tripod *i*, with 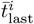 and 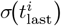 the mean and standard deviation, respectively; *T* ^∗^ = 0.25 *s* is the target gait period, so *T* ^∗^*/*2 is the target inter-tripod delay.

Double-tripod leg coordination was achieved via the contact reward term, which imposes an in-phase constraint within each tripod by minimizing the within-tripod variation in contact time, and sets a target contact delay between tripods. The orientation reward term minimizes the body’s roll, pitch, and yaw, and the antisymmetry term imposes equal-magnitude, opposite-sign angles in each pair of contralateral joints. The survival constant reward, together with early termination, encourages longer episodes.

The state of the model (joint positions and velocities and body position and orientation) was randomized at the beginning of each episode using uniformly-distributed random noise. Episodes lasted eight times the target period (unless early termination was triggered). All reward weights and early termination thresholds were fixed and identical across species, with the exception of the maximum *z*-position, which was set proportionally to each model’s height.

Optimization. We used Proximal Policy Optimization (PPO) with identical hyperparameters for both species. During optimization, reward terms act as soft constraints: the agent may trade small deviations from the target period or stance duration for greater forward velocity or stability, as reflected in the overall return. The optimization time step was *dt*_*opt*_ = 0.005 and the physics simulation time step was *dt*_*phy*_ = 0.001. The PPO agent was trained for a total of 12M time steps with 16 parallel environments.

### Extraction of locomotion parameters

Whole-body speed was defined as the change in body position in the x-axis per unit time. This was then averaged across all frames of each simulation episode, and then averaged across episodes.

The main body axis was defined as the vector from the abdomen center to the thorax center. Tip position of leg *i, d*_*i*_(*t*), was defined as the shortest distance of the tarsus segment from the secondary body axis (perpendicular to the main body axis). Using the maxima and minima in the graph of *d*_*i*_(*t*), kinematics-based stances were defined as the time intervals from the anterior-most (AEP) to the posterior-most (PEP) leg tip positions, and swings were defined as the remaining time intervals (Fig. 2C-D).

Leg angle was defined as the angle between the vector connecting the leg base to its tip, and the main body axis. To obtain the average leg angle, the circular mean was first used to average leg angles over time and then across episodes. Similarly, the circular standard deviation was used to obtain the standard deviation of leg angles over time, and then averaged across episodes using a regular mean (Fig. 2G-H).

Joint torque was defined as the controller’s action associated the joint. The torques of all joints were averaged and divided by the total body mass. This normalization makes sense because the gear ratio in all motors was scaled proportionally to the whole body masses of the two species.

The distance of leg trajectories from the thorax was computed by measuring the distance of each leg trajectory’s mean coordinate from the thorax center point.

The area of leg trajectories was measured as the area of their bounding envelopes. The bounding envelope was found using the “alphashape” Python library, which computes the bounding polygon (concave hull) containing a set of points. The shrinking factor (alpha) controls how tight the bounding polygon is, and was set to 0.5. Since the area measurement is sensitive to single points, we also computed the spread of each leg trajectory, defined as the mean distance of all trajectory coordinates from the mean trajectory coordinate.

Trajectory orientation angle was computed using the SVD method and by calculating the angle of the first principal component in [0, 90^*◦*^].

## 4 Discussion

Here we show that biologically constrained morphology and passive joint dynamics shape insect locomotion in simulation. We integrated morphometric measurements and passive femur–tibia (FT) dynamics into physics-based models and tested their consequences for learned walking. Our three key findings, elaborated below, include: (i) species-specific limb hypertrophies—enlarged mole-cricket forelegs and enlarged locust hind legs—yet conserved whole-body balance, with overall center-of-mass position maintained; (ii) convergent coordination with divergent kinematics/effort—under identical objectives both species learned stable double-tripod walking, attaining similar forward speeds but exhibiting species-specific differences in segment kinematics and normalized energetic cost; and (iii) material influence of passive properties—modifying only the passive parameters of the two hind FT joints produced a marked improvement: higher forward speed at lower joint torque. Together, these results demonstrate that adding measured morphometrics and passive mechanics narrows the parameter space, yielding more realistic behavior and stronger predictive leverage for both biological inference and bio-inspired design.

Under a shared objective that rewards forward advance and ground-contact timing, double-tripod coordination emerged in both species without an explicit phase oscillator, consistent with stick-insect models in which local sensory rules (e.g., ground contact/load and position feedback) suffice for interleg coordination [20, 21], indicating that ground-contact information can serve as a sufficient coordination signal in closed loop. At a higher level, the two insects converged on the same coordination pattern but diverged in how legs occupy and use workspace: across all limbs, locust trajectories span larger envelopes and lie farther from the body, whereas mole-cricket trajectories are tighter and more body-proximal, consistent with life underground and navigation in narrow burrows (Fig. 2A-B, K-M). Trajectory orientation differences map onto limb roles: a similar hind-leg orientation suggests similar propulsion constraints; locust middle-legs adopt a more body-axis-aligned posture consistent with their posterior placement; and mole-cricket forelegs are more body-axis-aligned, however they have near-circular workspaces, making principal-axis angle a weak descriptor for those digging-adapted limbs (Fig. 2A-B, N). These distinct features are consistent with morphological specialization—digging-adapted forelimbs in mole crickets versus jumping-biased posterior limbs in locusts—shaping which parts of the limb workspace are recruited during walking. Together, these findings indicate that high-level gait organization can converge across species, while species-specific morphology and joint workspaces sculpt where in space legs operate and how much workspace they recruit—yielding comparable coordination realized through distinct kinematic solutions.

Both models achieved the same forward speed, but the mole-cricket controller did so with a lower energetic cost (relative to its total body weight) (Fig. 2I-J), indicating a more economical solution under its morphology. Overall, imposing biologically grounded mechanical constraints on locomotion exposes meaningful, morphology-dependent, converging and diverging gait parameters, and predicts the energetic cost associated with body morphology during locomotion. Future research may test potential species-specific contributors, such as joint-range/posture defaults and differences in mass distribution or motor torque scaling.

Passive joint mechanics set fundamental limits on stability, timing, and energetic cost in legged systems; we therefore quantified and modeled the locust hind femur-tibia (FT) joint as a first step toward anatomically grounded constraints for whole-body simulation. The experiments revealed three robust features: a biphasic return (fast then slow), history-dependent resting angles with a flexion-extension dead-zone, and a small displacement-dependent rebound between phases (Fig. 3D-G). The transient rebound between the fast and slow return phases suggests an elastic recoil component beyond simple linear damping. Ahn and Full identified distinct “motor” and “brake” roles of extensor muscles that could contribute to rapid deceleration [7], but resilin-rich cuticular structures and tendon-like apodemes are likely key elastic elements. Targeted material tests on isolated joint cuticle—similar to approaches used by Dudek and Full to measure stiffness and damping in insect legs [18]—would quantify the nonlinear elasticity required to reproduce the observed rebound.

Our biologically-constrained joint model reproduced the fast-slow return and the hysteresis/dead-zone behavior, thus explaining much of the observed relaxation dynamics within a compact, mechanistic parameterization. However, the model did *not* reproduce the transient rebound, pointing to missing ingredients such as nonlinear stiffness, state-dependent damping, geometric contact effects, or additional elastic elements (e.g., cuticular springs) that merit targeted tests. Crucially, by modifying only the hind FT joints (and proportionally scaling their associated actuation to preserve feasible torques), holding morphology and all training settings fixed, the agent achieved higher forward speed at a lower energetic cost (Fig. 3K-L). Consistent with this economy shift, the constrained model reorganized limb workspaces—hind strokes moved closer to the body, swept larger envelopes, and aligned more with the body axis, while fore/middle strokes became more lateral—kinematic signatures that channel stance forces more effectively into forward propulsion (Fig. 3MS). This demonstrates that realistic passive constraints can improve locomotor economy and performance by shaping the dynamics the controller exploits, and underscores the value of measurement-informed models for predictive simulation. Moving forward, in situ force-angle and torque-transient measurements at the FT joint (cf. [18]), and microscopy-based identification of putative elastic tissues can disambiguate mechanisms. Critically, extending the same measurement-modeling pipeline to all major joints in both species will clarify how passive properties vary across the limb chain and how they shape gait stability, coordination, and energetics at the whole-body level.

Finally, our anatomically grounded simulation framework offers a path toward bio-inspired robotics by illustrating how real-world constraints improve both realism and predictive power. By systematically varying morphological and passive-mechanical parameters in silico, one can identify design principles that optimize stability and energy efficiency for different locomotor tasks. Building physical prototypes that implement these principles will close the loop between biological insight and engineered systems. Extending morphometric mapping and passive-mechanics measurements to all major joints in both species—and testing how these empirically derived constraints shape gait performance in simulation and in soft-robotic analogs—will broaden comparative inferences and sharpen model predictions.

## Declarations

### Ethics approval and consent to participate

Not applicable.

### Consent for publication

Not applicable.

### Availability of data and material

All the data are shown in the figures of this paper.

### Author contribution

Omer Yuval, Elad Ozeri, Avi Amir, and Amir Ayali conceived the study and planned the experiments. Avi Amir collected and maintained the animals. Elad Ozeri performed the in vivo joint experiments, including video acquisition, and analyzed the recordings. Omer Yuval and Elad Ozeri developed the biologically-constrained joint model. Omer Yuval developed the biomechanical insect models and the reinforcement learning control framework, analyzed the simulation data, and wrote the original draft. Amir Ayali acquired funding and supervised the project. All authors reviewed and approved the final manuscript.

